# A fast nociceptive subsystem mediating rapid reflexive behavior but not affective pain

**DOI:** 10.1101/2025.03.17.643731

**Authors:** Chwen-Yu Chen, Leandro Flores do Nascimento, Felipe Meira de Faria, Gabriela Carballo, Lech Kaczmarczyk, Jonas Broman, Saad Nagi, Håkan Olausson, Walker S Jackson, Marcin Szczot, Max Larsson

## Abstract

Spinal nociceptive withdrawal reflexes are widely believed to rely on unmyelinated and thinly myelinated nociceptive fibers that also signal affective and motivational aspects of pain. Here we discover a population of myelinated mechanoreceptive nociceptor that forms free nerve endings as well as circumferential endings around hair follicles, and exclusively terminate in the deep spinal dorsal horn. Optogenetic activation of these fibers triggers rapid withdrawal reflexes that are precise and selective for the targeted limb, while silencing increases the threshold of mechanical nociceptive withdrawal reflexes. By contrast, optogenetic stimulation of the fibers is not associated with place aversion nor with changes in facial expression. Thus, we conclude that this nerve fiber population is uniquely positioned to rapidly respond to mechanical threats via selective withdrawal of the targeted body part, whereas other fast and slow nociceptive pathways are required for affective-motivational aspects of pain.

## Introduction

One of the most critical tasks of the nervous system is to alert the organism to impending threats to body integrity by stimuli emanating from the external environment. Many mechanical stimuli may adversely affect the tissue much more rapidly than either heat, cold or chemical stimuli, and very fast nociceptive withdrawal reflexes (NWRs) have evolved to specifically counter imminent mechanical threats. Notably, while the term nociception encompasses both signals that mediate NWRs as well as discriminative and affective/motivational aspects of pain, it is widely thought that specific nociceptive primary afferent fibers generally mediate both NWRs and pain. Moreover, it has been suggested that mechanical stimulation of nociceptors alone triggers exaggerated, suboptimal reflexive behavior and that concurrent activation of low-threshold tactile afferent fibers serve to sculpt the NWR to produce well-coordinated, appropriate motor action (1).

Since the existence of specialized peripheral nociceptors was confirmed (2), extensive efforts have been devoted to identifying neurochemical and physiological markers for nociceptor subgroups. With the advent of genetic tools for manipulation of select fiber populations, it has become possible to interrogate functional differences between select nociceptor populations. For instance, it was found that two populations of unmyelinated C fibers induce distinct behavior on activation – those carrying the heat-activated receptor TRPV1 induced licking behavior and place aversion, whereas C fibers with the MrgprD receptor induced NWR but no place aversion (3-5). However, C fiber activation cannot explain the generation of rapid mechanical NWRs since the latency to paw withdrawal mediated by MrgprD fibers is several hundred milliseconds, a magnitude too slow to account for the rapid mechanically evoked NWR (6). In the present study we characterized a small subpopulation of myelinated nociceptor in the mouse, finding that these fibers are sufficient and necessary for normal rapid NWR induction, but appear to not trigger affective pain. Thus, we demonstrate a dissociation between rapid nociception and pain already at the first step of somatosensation.

## Results and discussion

During a search for new populations of myelinated A fiber nociceptors, we noted in *in situ* hybridization experiments that transcripts for *Ntrk3*, encoding the TrkC receptor, and *Scn10a*, which encodes the voltage-gated Na^+^ channel Na_V_1.8 that is mostly restricted to nociceptors (7), co-localized in a population of medium-to-large sized neurons in dorsal root ganglia (DRGs) (Fig. 1a). To characterize this population we employed an intersectional genetic targeting approach, crossing a newly generated Na_V_1.8^FlpO^ mouse line (8) with TrkC^CreERT2^ mice (9); this cross is hereafter called Na_V_1.8^TrkC^. In this manner, targeting of TrkC^+^ A fiber low-threshold mechanoreceptors (A-LTMRs) that lack Na_V_1.8 (9, 10) could be avoided. In Na_V_1.8^TrkC^ mice crossed with Ai65 mice that conditionally express tdTomato, after tamoxifen administration at 6-10 weeks of age, recombined neurons accounted for 2 % of DRG neurons at the lumbar L3-4 level. These neurons were of medium-to-large diameter and did not co-localize with the common nociceptive markers CGRP, TRPV1 or IB_4_ binding (Fig. 1c-e). Moreover, tdTomato^+^ neurons expressed the myelinated neuron marker NFH, and in the sciatic nerve, Na_V_1.8^TrkC^ fibers were generally thick and thickly myelinated as compared to CGRP^+^ axons (Fig. 1f, g). In glabrous skin, Na_V_1.8^TrkC^ fibers formed free nerve endings, and in hairy back skin, in addition to epidermal free nerve endings, also circumferential nerve endings around some hair follicles (Fig. 2a,b).

**Figure 1.**
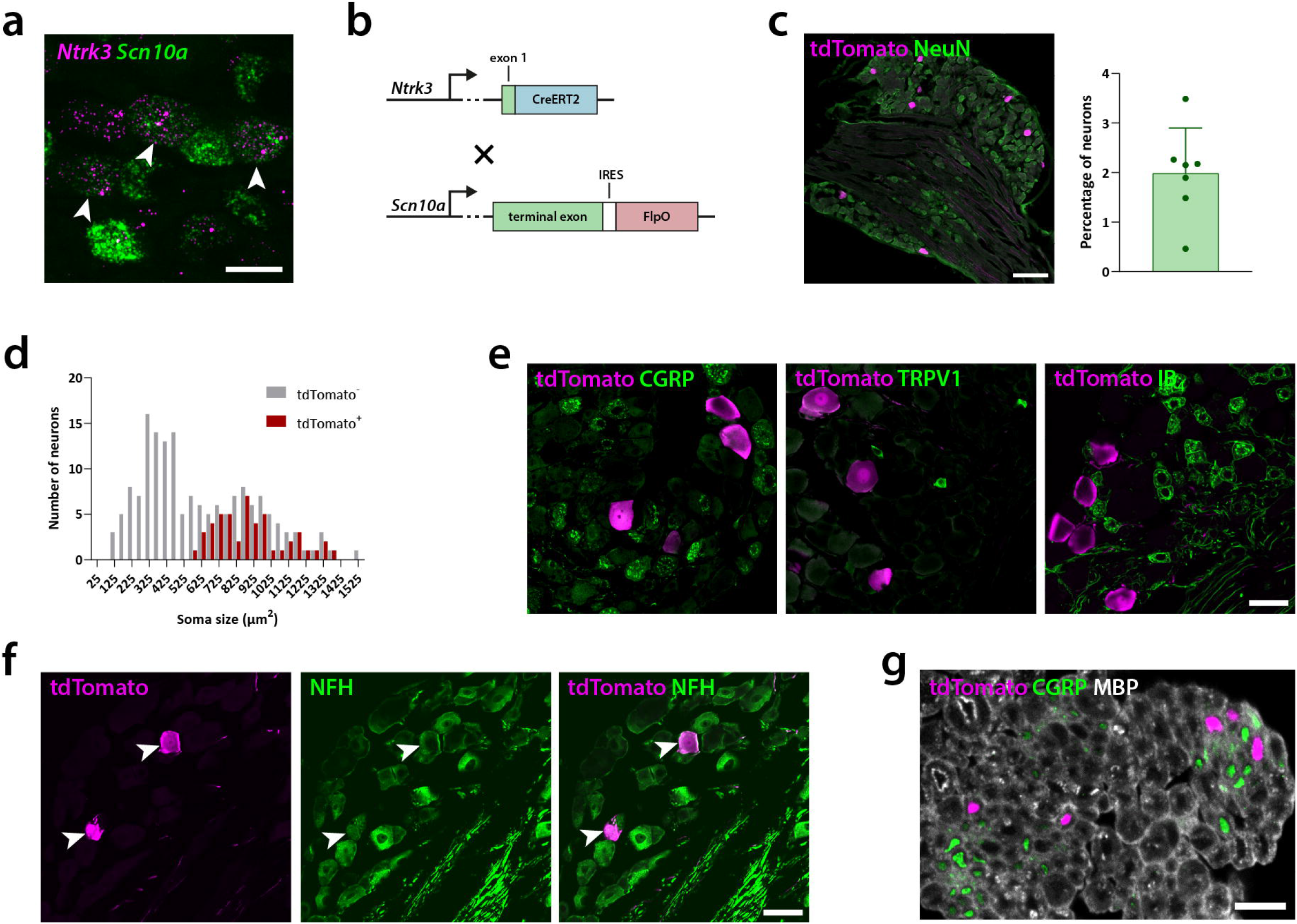
Genetic targeting of Na_V_1.8^TrkC^ DRG neurons. **a**, in situ hybridization of *Ntrk3* (encoding TrkC) and *Scn10a* (encoding Na_V_1.8) transcripts in a lumbar DRG of a C57Bl/6 mouse shows cells co-expressing these genes. Arrowheads indicate cells co-localizing *Ntrk3* and *Scn10a* mRNA. Scale bar, 50 µm. **b**, schematic view of the intersectional approach for targeting Na_V_1.8^TrkC^ neurons using TrkC^CreERT2^ and Na_V_1.8^FlpO^ mice. **c**, recombined cells in L3-L4 DRGs in TrkC^CreERT2^;Na_V_1.8^FlpO^;Ai65 mice. tdTomato^+^ cells are ∼2 % of NeuN^+^ DRG neurons. Scale bar, 200 µm. **d**, soma size of tdTomato^+^ DRG neurons as compared to tdTomato-neurons. **e**, co-localization of tdTomato with nociceptive markers CGRP and TRPV1 and IB_4_ binding. No co-localization was found for either marker. Scale bar, 50 µm, valid for all panels. **f**, co-localization with the myelinated neuron marker neurofilament heavy chain (NFH). Arrowheads indicate tdTomato^+^ neurons that also are immunoreactive for NFH. Scale bar, 50 µm. **g**, a section of sciatic nerve co-stained for tdTomato, CGRP and the myelin marker myelin basic protein (MBP). Scale bar, 10 µm.

**Figure 2.**
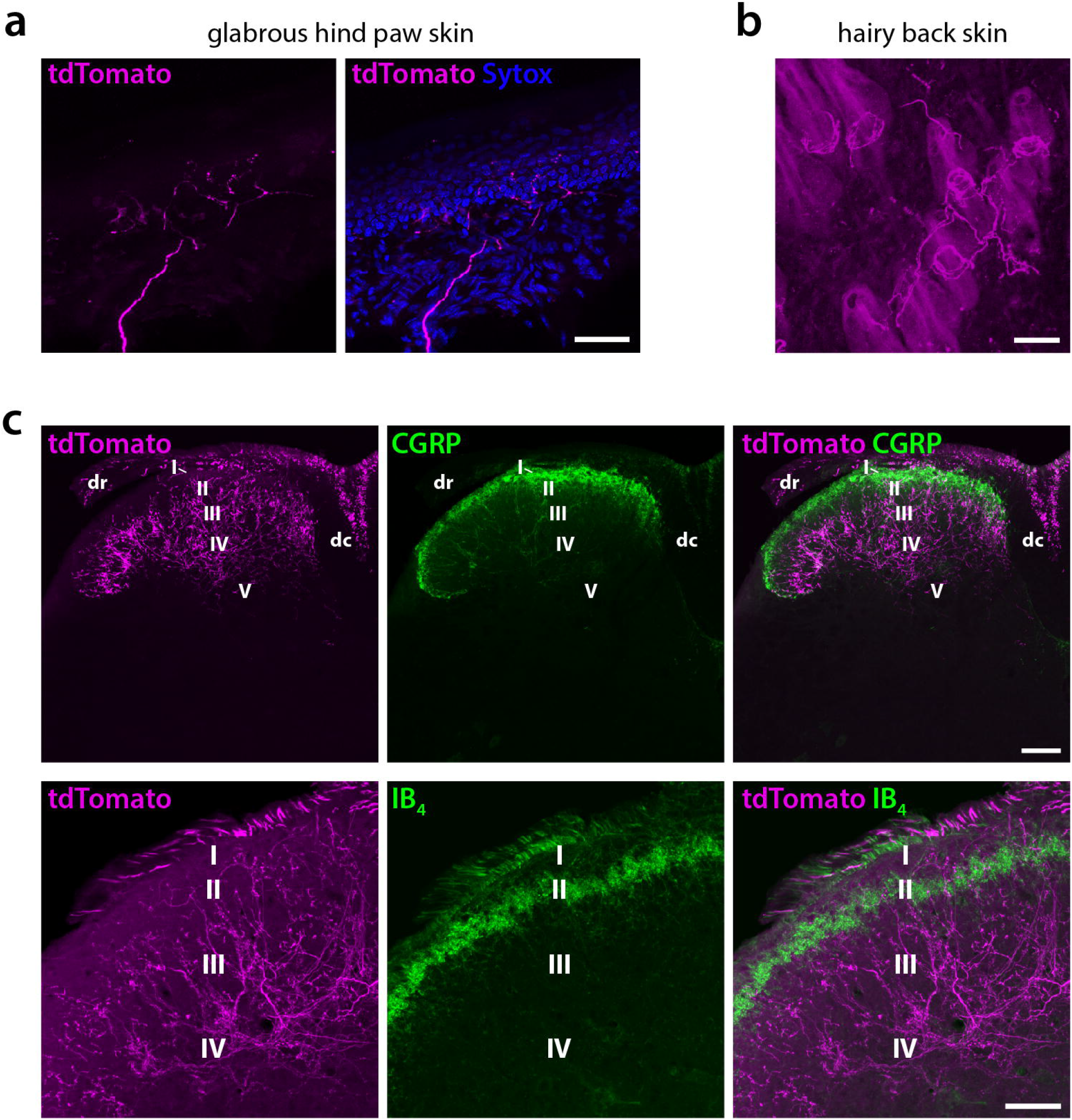
Cutaneous and central processes of Na_V_1.8^TrkC^ neurons. **a**, a tdTomato^+^ fiber establishing epidermal free nerve endings in glabrous hind paw skin from a TrkC^CreERT2^;Na_V_1.8^FlpO^;Ai65 mouse. Scale bar, 50 µm. **b**, tdTomato^+^ fibers forming circumferential nerve endings around several hair follicles in back skin. Note that epidermal free nerve endings are not discernible in this view. Scale bar, 50 µm. **c**, central tdTomato^+^ processes in the lumbar spinal cord. Processes were abundant in lamina II_inner_-IV, but essentially absent from lamina I and V. Top panels show the distribution pattern of tdTomato^+^ processes compared to CGRP^+^ fibers, which are abundant in lamina I and II_outer_. Note the minimal overlap between CGRP and tdTomato. Scale bar, 100 µm. Bottom panels show a higher magnification view of a mid-lateral portion of the dorsal horn co-labeled for tdTomato and IB_4_, which in mice mark the middle third of lamina II. Most tdTomato^+^ processes were ventral to the IB_4_ plexus although a few processes entered this region. Scale bar, 50 µm.

This structurally distinct innervation of epidermis and hair follicles could be because of a remaining heterogeneity within the Na_V_1.8^TrkC^ population, or to dual innervation by single neurons. Remarkably, in the spinal cord, Na_V_1.8^TrkC^ fibers were essentially absent from both laminae I and V, which are conventially considered the major central targets of A-nociceptors (11, 12, but see 13). Instead, Na_V_1.8^TrkC^ fibers populated laminae III-IV, which is generally considered the LTMR receptive field of the spinal cord (Fig. 2c). Some Na_V_1.8^TrkC^ fibers entered the mid- and inner part of lamina II, which also receives primary afferent input from Aδ hair afferents and C-LTMRs (14). Still, in accordance with the low proportion of Na_V_1.8^TrkC^ cells in the DRG, fibers in the dorsal horn were sparse when compared with CGRP^+^ nociceptors and IB_4_ binding non-peptidergic C nociceptors.

To determine adequate stimuli for Na_V_1.8^TrkC^ fibers, we performed *in vivo* Ca^2+^ imaging in L3-L4 DRGs of Trkc^CreERT2^;NaV1.8^FlpO^;Ai195 mice that expressed GCaMP7f in a Cre/Flp-dependent manner. Na_V_1.8^TrkC^ cells with receptive field in the plantar hind paw were rare, in general only 1-2 such cells were visible at the surface of the ganglion, and in some DRGs no cells were found. Na_V_1.8^TrkC^ cells with plantar receptive field were reliably activated by pin prick, sometimes by sandpaper moving horizontally across the skin at low downward force, but not by a velvet-covered flat block applied similarly to the sandpaper (Fig. 3a). Thus, Na_V_1.8^TrkC^ cells with input from hind paw glabrous skin are moderate-to-high threshold mechanoreceptors.

**Figure 3.**
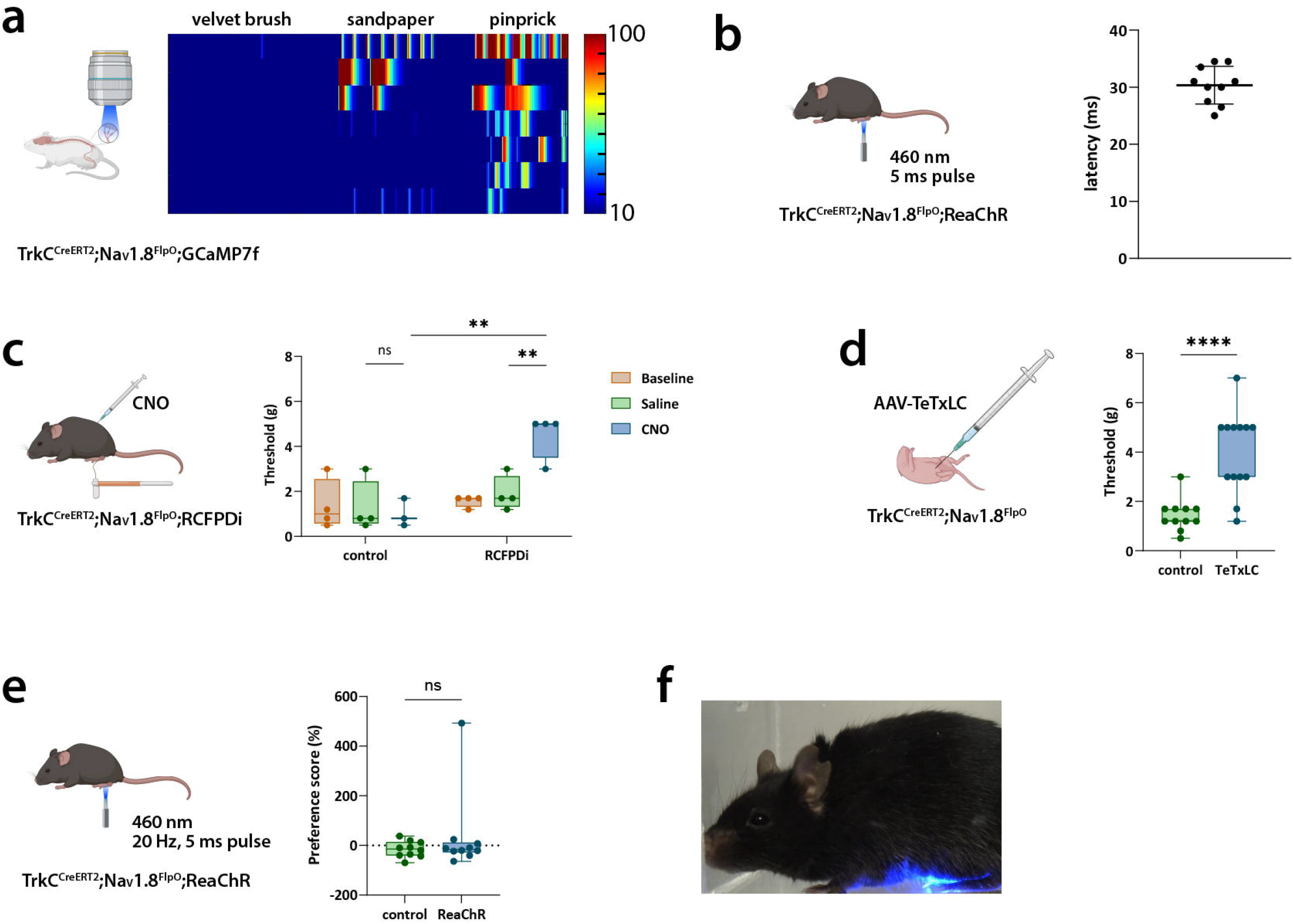
Functional characterization of Na_V_1.8^TrkC^ neurons. **a**, *In vivo* Ca^2+^ imaging in L3-L4 DRG of TrkC^CreERT2^;Na_V_1.8^FlpO^;GCaMP7f mice. Neurons responded to pin prick, sometimes sandpaper but not to velvet brush applied to glabrous hind paw skin. **b**, withdrawal latencies to optogenetic stimulation of Na_V_1.8^TrkC^ fibers in plantar hind paw. **c**, mechanical threshold to paw withdrawal was increased by chemogenetic inhibition of Na_V_1.8^TrkC^ fibers. **d**, mechanical threshold to paw withdrawal was higher in mice where synaptic transmission from Na_V_1.8^TrkC^ fibers was abolished using TeTxLC expressed via pup injections of AAV encoding this toxin in a Cre/Flp-dependent manner. **e**, realtime place preference assay showed no aversion to optogenetic stimulation of Na_V_1.8^TrkC^ fibers in the hind paw. **f**, no changes in facial expression indicative of affective pain were observed during optogenetic stimulation of hind paw Na_V_1.8^TrkC^ fibers.

In mice expressing the channelrhodopsin ReaChR in Na_V_1.8^TrkC^ fibers, a single light pulse (595 nm, 5 ms) applied to the plantar hind paw robustly produced a withdrawal of the paw (Fig. 3b). The movement of the paw was initiated with a latency of 30.4 ± 3.2 ms, and was restricted to the stimulated paw; no jumping or lifting of the contralateral paw occurred. Chemogenetic inhibition of Na_V_1.8^TrkC^ fibers using the DREADD hM4D_i_ in TrkC^CreERT2^;Na_V_1.8^FlpO^;RCFPDi mice resulted in a marked increase in mechanical threshold to paw withdrawal using von Frey filaments after i.p. injection of clozapine-*N*-oxide (CNO) compared to littermate control mice and after saline injection (Fig. 3c). In separate experiments, we then silenced synaptic transmission from Na_V_1.8^TrkC^ fibers using Cre/Flp-dependent expression of tetanus toxin light chain (TeTxLC), via neonatal i.p. injections of TeTxLC encoding adeno-associated virus (AAV). This resulted in a similar phenotype to that after chemogenetic inhibition, increasing the mechanical threshold to paw withdrawal (Fig. 3d). We conclude that Na_V_1.8^TrkC^ fibers are sufficient and essential for normal mechanically evoked rapid withdrawal reflexes.

We noted that optogenetic stimulation of Na_V_1.8^TrkC^ fibers in the hind paw, while evoking a robust paw withdrawal, did not appear to produce any overt discomfort in the animal, and rarely generated any attention of the animal to the stimulated paw. Indeed, in a realtime place preference assay, there was no aversion of the mice to optogenetic stimulation of the hind paw (Fig. 3e). Moreover, during a 3-min period of continuous optogenetic stimulation at 20 Hz, no overt changes in facial expression were observed, and specifically no changes consistent with those found when applying other types of nociceptive stimulation, such as orbital tightening or backward orientation of the ears (15, 16)(Fig. 3f). Thus, we found no indication that activation of Na_V_1.8^TrkC^ fibers by itself triggers activation of nociceptive pathways that induce affective pain.

In conclusion, here we report that a new population of cutaneous A-mechanonociceptor is instrumental for the initiation of NWRs by mechanical stimuli, but is not capable of driving aversive behavior linked to affective-motivational facets of pain. Our observation that selective stimulation of these fibers triggers well-coordinated, somato- and myotopically appropriate NWRs suggests that simultaneous activation of low-threshold Aβ mechanoreceptors is not necessary to prevent the generation of exaggerated reflexive movements (1). Because C fiber nociceptors are too slow to account for rapid mechanically evoked NWRs, we posit that Na_V_1.8^TrkC^ fibers are a major driver of such protective reflexes.

## Methods

### Animals

The following mouse lines were used: C57BL/6JRj, Na_V_1.8^FlpO^ (8), TrkC^CreERT2^ (Jax #030291) (9), Ai65 (RCFL-tdT; Jax# 021875), LSL_FSF_ReaChR (Jax# 024846), RCFPDi (Jax #029040) (17). The animals were housed in groups of 2-4 same-sex individuals in NexGen IVC system Mouse 500 home cages in a 20±2°C environment, under a 12L:12D light cycle with *ad libitum* access to food and water. The nocturnal phase lasted from 19.00 to 7.00. All animal experiments were approved by the Animal Ethics Committee at Linköping University (permit no. 2439-2021) and performed in accordance with the EU Directive 2010/63/EU.

### Tamoxifen administration

Tamoxifen (Sigma-Aldrich T5648) was dissolved in corn oil at 20 mg/mL concentration overnight at 37°C. Mice at 6-10 weeks of age received one tamoxifen injection (100 µL, i.p.). The mice were subjected to further experiments not earlier than two weeks after tamoxifen injections.

### Tissue preparation

Mice of either sex were anesthetized with sodium pentobarbital (100 mg/kg) and subjected to transcardial perfusion using 5 mL phosphate buffer (PB, 0.1 M, pH 7.4) followed by 50 mL of 4 % paraformaldehyde. Tissue (brain, spinal cord, dorsal root ganglia, sciatic nerves, back skin and hind paw skin) was harvested, post-fixed in fixative at 4°C overnight and stored in 1/10 fixative at 4°C before further processing.

### Immunofluorescence

Tissue specimens from TrkC^CreERT2^;Na_V_1.8^FlpO^;Ai65 mice (2 females, 3 males, 10-13 weeks old) and littermate controls (1 female, 1 male; 11 and 13 weeks old) were cryoprotected in 30 % sucrose and embedded in OCT. Sections were cut at 15-50 µm thickness in a cryostat and placed on slides. The sections were incubated in phosphate-buffered saline (PBS) with 3% normal goat serum, 0.5 % bovine serum albumin and 0.5 % Triton X-100 (blocking solution), followed by incubation in primary antibodies (see Table 1) diluted in blocking solution at room temperature overnight. After rinsing, the sections were incubated in blocking solution containing secondary antibodies at a dilution of 1:500. To detect binding sites for isolectin B_4_ (IB_4_), biotinylated IB_4_ (dilution 1:2000) in conjunction with streptavidin-Alexa 488 (dilution 1:500; Life Technologies) was used. In skin sections, cellular nuclei were labeled in a separate step using DAPI (1:1000; Life Technologies) or SYTOX Deep Red (1:2000). Sections were coverslipped with Prolong Glass or SlowFade Diamond (Life Technologies). Sections were imaged using a Zeiss LSM800 confocal microscope, or a Leica Stellaris 5 confocal microscope.

**Table 1.** Primary antibodies.

| Antigen | Host, isotype | Supplier | Cat # | Dilution |
| --- | --- | --- | --- | --- |
| CGRP | Guinea pig | Synaptic Systems | 414 004 | 1:1000 |
| NFH | Chicken | Thermo Fisher Scientific | PA1-10002 | 1:1000 |
| NeuN | Mouse, IgG<sub>1</sub> | Merck Millipore | MAB377 | 1:1000 |
| RFP (tdTomato) | Chicken | Synaptic Systems | 409 006 | 1:2000 |
| RFP (tdTomato) | Rabbit | Rockland | 600-401-379 | 1:1000 |
| TRPV1 | Rabbit | Synaptic Systems | 444 033 | 1:1000 |

### In situ hybridization

Lumbar DRGs from adult C57BL/6JRj mice of either sex were cryoprotected in 30 % sucrose, and embedded in OCT. Sections were cut in a cryostat at 15 µm thickness and placed on SuperFrost slides. *In situ* hybridization of *Ntrk3* and *Scn10* was performed using the manual RNAscope (Advanced Cell Diagnostics) procedure according to the standard protocol.

### *In vivo* imaging

TrkC^CreERT2^;Na_V_1.8^FlpO^ animals were crossed with Ai195 mice to generate TrkC^CreERT2/wt^;Na_V_1.8^FlpO/wt^;GCaMP7f^+/wt^ mice. Tamoxifen was administered as above. Adult mice of both sexes were used. For the imaging procedure, the mouse was anesthetized with isoflurane (4% induction, 1.5% maintenance) and transferred to a custom surgical platform equipped with a heating pad for body temperature maintenance. The lumbar spine was surgically exposed, stabilized with a spinal clamp (Narishige STS-A), and a dental drill used to remove bone in order to gain visual access to the L3-4 DRGs. Following surgery, the animal was transferred to the stage of a custom tilting light microscope (Thorlabs Cerna) equipped with a 4x/NA 0.28 dry objective (Thorlabs). Fluorescence images were acquired for 40 second epochs at 5 Hz with an sCMOS camera (Sona 4.2, Andor) using a standard green fluorescent protein filter cube. Stimuli were applied to the plantar aspect of either hind paw multiple times for the duration of the recording. The velvet brush stimulus was applied using a custom brush consisting of a handle ending in a head with a raised 6×8 mm flat surface, onto which a piece of velvet fabric of the same size was glued. The stimulus was applied by manually moving the velvet surface in a slow (∼1 cm/s) proximodistal motion along the glabrous skin from the heel to the tips of the digits, with little downward force. Sandpaper stimulus was applied as the velvet brush, except that a coarse (P60 grit) sandpaper was substituted for the velvet fabric. The pin prick stimulus was applied using a 3D-printed rigid resin block with a rectangular grid of conical pins, each ending in a tip of 200 µm diameter; the grid consisted of 30 pins arranged in an equidistant rectangular pattern within an area of 20×5 mm. This pin block was evenly applied with about 300 mN force onto the plantar hind paw skin (thus yielding ∼10 mN of force per pin); downward force was limited by placing the paw on a mount that was 3D printed from flexible resin (Liqcreate Flexible-X) and constructed so as to bend above a certain downward force.

Analysis of Ca^2+^ imaging was performed as previously described (18). Regions of interest (ROI) outlining responding cells were drawn in FIJI/ImageJ using the Cell Wand Tool plugin and relative change of GCaMP7f fluorescence was calculated as percent ΔF/F. Background signal (*e*.*g*., from out-of-focus tissue and neighboring cells), was removed by subtracting the fluorescence of a donut-shaped area surrounding each ROI using a custom-written MATLAB script. Overlapping ROIs and rare spontaneously active cells were not included in the analysis. Imaging episodes were concatenated for visualization as activity heatmaps where final ΔF/F was shown from 10 to 100%.

### Behavioral assays

To assess latency of optogenetically induced withdrawal reflexes, TrkC^CreERT2^;Na_V_1.8^FlpO^;ReaChR mice were singly placed in a transparent Plexiglass cubicle (9×5×5 cm) with a 3 mm thick floor, and left to habituate for 10 min. A fiberoptic cable (0.63 NA, 960 µm) attached to a 460 nm LED (Prizmatix) was used to apply a single pulse of light (5 ms) to the plantar hind paw through the floor. The latency to paw withdrawal was measured from the start of the pulse to the start of paw movement in videographs recorded at 1000 fps. Each hind paw was stimulated once and the mean of the two latencies calculated.

For realtime place preference, a Plexiglass apparatus composed of two 175×90 mm chambers connected via a 50 mm opening was used. Tests were recorded using a web camera (Logitech C920) and AnyMAZE software. On Day 1, TrkC^CreERT2^;Na_V_1.8^FlpO^;ReaChR and littermate mice were placed in the apparatus and allowed to explore freely for 15 min (pre-test). On Day 2, the mice were again placed in the apparatus for 15 min; one hind paw was continuously tracked with the tip of a fiberoptic cable (0.63 NA, 960 µm, attached to a Prizmatix 460 nm LED) manually placed just below the plantar hind paw through the transparent 5 mm thick Plexiglass floor. When the mouse entered the chamber initially preferred during pre-test, optogenetic stimulation was switched on, controlled via AnyMAZE software, and remained on until the mouse exited the chamber. Preference score was calculated as 100*(t_test_ – t_pre_) / t_pre_, where t_test_ is time spent in light-on chamber during test and t_pre_ is time spent in the same chamber during pre-test.

Genotypes were unknown to the experimenter at the time of testing.

For von Frey assays during chemogenetic inhibition, TrkC^CreERT2^;Na_V_1.8^FlpO^;RCFPDi animals and control littermates were on Day 1 placed in a cubicle (9×5×5 cm) with a mesh floor. After 30 min habituation, the baseline mechanical threshold to hind paw withdrawal was assessed using von Frey filaments and the simple up-down protocol (ref). On Days 2 and 3, the mice were injected with either saline or CNO (1 mg/kg i.p.), habituated for 30 min and assessed for von Frey threshold. The order of saline versus CNO injection was randomized for each mouse, and the experimenter was blind to both treatment and genotype.

For von Frey assays in mice where synaptic transmission from Na_V_1.8^TrkC^ fibers had been abolished using TeTxLC, TrkC^CreERT2^;NaV1.8^FlpO^ pups (P1-P5) were briefly anesthetized with isoflurane and injected i.p. with AAV9.Con/Fon-TeTxLC (1 µL of 8.2×10^12^ vg/mL solution diluted 1:10 in saline, for a final injected volume of 10 µL). Each pup received two injections, either at P2 and P4, or P3 and P5. The TeTxLC plasmid was constructed by Dr Hendrik Wildner, University of Zürich, and the AAVs produced by the University of Zürich Viral Vector Facility. After tamoxifen injections at 6-10 weeks age, the mice were subjected to von Frey assays as above.

TrkC^CreERT2/wt^;Na_V_1.8^FlpO/wt^ and TrkC^wt/wt^;Na_V_1.8^FlpO/wt^ littermates were used; the experimenter was blind with respect to genotype.

For the facial expression assay, TrkC^CreERT2^;Na_V_1.8^FlpO^;ReaChR mice and littermate controls were placed in a cubicle (9×5×5 cm) similar to the withdrawal latency test above. Videography of the lateral view of the cubicle was recorded using a color FLIR camera (Blackfly BFS-U3-23S3C-C) with a 1:1.8/4 mm Basler lens (C125-0418-5M) and the Open Broadcaster Software 27.0.1 (https://obsproject.com/). Optogenetic stimulation was applied to the plantar hind paw through the floor via a fiberoptic cable (0.63 NA, 960 µm core) connected to a 460 nm LED (Prizmatix). After 10 min of habituation, baseline was recorded for 3 min, after which a hind paw was optogenetically stimulated (20 Hz, 5 ms pulses) for 3 min. If the targeted hind paw was moved out of reach for continued stimulation, the tip of the fiberoptic cable was repositioned to the other hind paw. After the stimulation, a 3 min recovery phase was also recorded. The experimenter was blind to the genotype during testing.

## Statistical analysis

GraphPad Prism was used for all statistics. Data are presented as mean ± S.D. where applicable, or as median ± range.

## References

1. Arcourt, A., et al. (2017) Neuron 93, 179–93.

2. Burgess, P.R. and Perl, E.R. (1967) J Physiol 190, 541–62.

3. Beaudry, H., et al. (2017) PAIN 158,

4. Warwick, C., et al. (2021) PAIN 162, 2120–31.

5. Huang, T., et al. (2019) Nature 565, 86–90.

6. Abdus-Saboor, I., et al. (2019) Cell Reports 28, 1623–34.e4.

7. Kupari, J. and Ernfors, P. (2023) Pain 164, 1245–57.

8. Chen, J.C.-Y., et al. (2025) 10.13140/RG.2.2.21502.91204.

9. Bai, L., et al. (2015) Cell 163, 1783–95.

10. Handler, A. and Ginty, D.D. (2021) Nat Rev Neurosci 22, 521–37.

11. Todd, A.J. (2010) Nat Rev Neurosci 11, 823–36.

12. Willis, W. and Coggeshall, R. (2004)

13. Boada, M.D. and Woodbury, C.J. (2008) J Neurosci 28, 2006–14. 14.

14. Li, L., et al. (2011) Cell 147, 1615–27.

15. Le Moëne, O. and Larsson, M. (2023) eNeuro 10, ENEURO.0349-22.2022.

16. Le Moëne, O. and Larsson, M. (2025) Affect Sci 6, 159–70. 17.

17. Ray, R.S., et al. (2011) Science 333, 637–42.

18. Ghitani, N., et al. (2017) Neuron 95, 944–54.e4.

